# Substitutions and codon usage in SARS-CoV-2 in mammals indicate natural selection and host adaptation

**DOI:** 10.1101/2021.04.04.438417

**Authors:** Zhixiong Lei, Dan Zhang, Ruiping Yang, Jian Li, Weixing Du, Yanqing Liu, Huabing Tan, Zhixin Liu, Long Liu

## Abstract

The outbreak of COVID-19, caused by severe acute respiratory syndrome coronavirus 2 (SARS-CoV-2) infection, rapidly spread to create a global pandemic and has continued to spread across hosts from humans to animals, transmitting particularly effectively in mink. How SARS-CoV-2 evolves in animals and humans and the differences in the separate evolutionary processes remain unclear. We analyzed the composition and codon usage bias of SARS-CoV-2 in infected humans and animals. Compared with other animals, SARS-CoV-2 in mink had the most substitutions. The substitutions of cytidine in SARS-CoV-2 in mink account for nearly 50% of the substitutions, while those in other animals represent only 30% of the substitutions. The incidence of adenine transversion in SARS-CoV-2 in other animals is threefold higher than that in mink-CoV (the SARS-CoV-2 virus in mink). A synonymous codon usage analysis showed that SARS-CoV-2 is optimized to adapt in the animals in which it is currently reported, and all the animals showed decreased adaptability relative to that of humans, except for mink. A binding affinity analysis indicated that the spike protein of the SARS-CoV-2 variant in mink showed a greater preference for binding with the mink receptor ACE2 than with the human receptor, especially as the mutation Y453F and F486L in mink-CoV lead to improvement of binding affinity for mink receptor. Our study focuses on the divergence of SARS-CoV-2 genome composition and codon usage in humans and animals, indicating possible natural selection and current host adaptation.

## Introduction

SARS-CoV-2 is a β-coronavirus that emerged in 2019 and spread worldwide, leading to an ongoing global pandemic [1,2]. As of February 19^th^ 2021, the number of infected cases reached 110 million, and more than 2.4 million deaths have occurred (Johns Hopkins University statistics; https://coronavirus.jhu.edu/map.html). SARS-CoV-2 has a single-stranded positive-sense RNA genome containing 29,903 nucleotides and consisting of 11 open reading frames (ORFs) encoding 27 proteins[3]. The S glycoprotein is a fusion viral protein that functions in recognition of the host receptor ACE2[4].

There is a broad host spectrum because SARS-CoV-2 binds a receptor common to humans and animals[5]. To date, the following animals have been reported to be susceptible to infection: cats, dogs, tigers, lions, ferrets, and mink [6–12]. SARS-CoV-2 infection of pets, including cats and dogs [8,10], was the earliest reported animal infections in the epidemic. Later, in a report on SARS-CoV-2 infection in tigers, lions, and human keepers in a New York zoo [11], epidemiologic and genomic data indicated human-to-animal transmission [13]. Other animals, including snow leopards and gorillas, tested positive for SARS-CoV-2 after showing signs of illness [14,15]. It is noteworthy that a study from The Netherlands reported the spread of SARS-CoV-2 from humans to mink and from mink back to humans in mink farms [16]. Eighty-eight mink and 18 staff members from sixteen mink farms were confirmed to be infected with SARS-CoV-2 as determined by sequence analysis. The adaptation of SARS-CoV-2 to bind the mink receptor and the viral evolution in the mink host are worthy of further study.

Codon usage bias refers to differences in the frequency of occurrence of synonymous codons during protein translation, which differs between hosts [17]. Viruses differ markedly in their specificity toward host organisms, and the analysis of the viral genome structure and composition contributes to the partial understanding of virus evolution and adaptation in the host [18]. Further exploration of the codon usage pattern of SARS-CoV-2 in different hosts, especially the codon architecture of the *Spike* gene, indicate host adaptation related to cross-species transmission.

Surveillance of the substitution and selection of the SARS-CoV-2 genome is important for the study of viral evolution and for tracking viral transmission. In particular, study of the *Spike* gene helps to evaluate the immunization effect of vaccinations and to adjust the vaccine design in a timely manner. This study focuses on the divergence of the SARS-CoV-2 genome composition and codon usage in human and animal hosts to investigate the natural selection that might play a role in virus evolution, adaptability, and transmission.

## Materials and Methods

### SARS-COV-2 sequences and data collection

A total 207 SARS-COV-2 genome sequences from humans, cats, dogs, tigers, lions, ferrets, and minks were used for the genetic analysis (The strains information was recorded in the Supplementary Table S1). All the genomic sequences selected by the hosts were obtained from the GAISD database (https://www.gisaid.org/). Isolate Wuhan/WIV04 was used as the reference strain.

### Evolutionary analysis

Thirty-nine SARS-COV-2 genomes were used for phylogenetic analysis. The evolutionary history was inferred by using the Maximum Likelihood method and Tamura-Nei model [19]. The tree with the highest log likelihood (−42442.50) is shown. The percentage of trees in which the associated taxa clustered together is shown next to the branches. Initial tree(s) for the heuristic search were obtained by applying the Neighbor-Joining method to a matrix of pairwise distances estimated using the Maximum Composite Likelihood (MCL) approach. A discrete Gamma distribution was used to model evolutionary rate differences among sites (5 categories (+G, parameter = 0.0500)). The tree is drawn to scale, with branch lengths measured in the number of substitutions per site. This analysis involved 39 nucleotide sequences. There was a total of 29903 positions in the final dataset. Evolutionary analyses were conducted in MEGA-X [20].

### Identification of mutations

The sequences were aligned using MEGA-X, and the single nucleotide polymorphisms were analyzed using the SNiPlay pipeline by uploading aligned Fasta format file (https://sniplay.southgreen.fr/cgi-bin/analysis_v3.cgi)[21]. All the sequences including coding regions, 5’UTR and 3’UTR were used for the analysis.

### Estimation of nonsynonymous and synonymous substitution rates

The number of nonsynonymous substitutions per synonymous site (dN) and the number of synonymous substitutions per nonsynonymous site (dS) for each coding site were calculated using the Nei–Gojobori method (Jukes–Cantor) in MEGA-X. The Datamonkey adaptive evolution server (http://www.datamonkey.org) was used to identify sites where only some of the branches have undergone selective pressure. The mixed-effects model of evolution (MEME) and fixed effects likelihood (FEL) approaches were used to infer the nonsynonymous and synonymous substitution rates.

### Codon usage analysis

The codon adaptation index (CAI) of a given coding sequence was calculated using R script [22]. A CAI analysis of those coding sequences from different hosts was performed using DAMBE 5.0 and the CAI [23,24]. The codon usage data of different hosts were retrieved from the codon usage database (http://www.kazusa.or.jp/codon/), and the relative synonymous codon usages (RSCUs) were analyzed using MEGA software.

### Spike protein sequence and structure reconstruction

The crystal structure of the SARS-CoV-2 receptor-binding domain (RBD) in complex with human ACE2 (PBD ID: 6M0J) was used for structural analysis. Structures of ACE2 and the viral spike from mink were constructed by the SWISS model server (https://swissmodel.expasy.org/). Comparisons of the predicted protein structures and pairwise comparisons were analyzed using PyMOL software.

### Molecular dynamics

For the binding free energy (*E*), we simulated the minimized annealing energy through molecular dynamics (MD) simulation in YASARA [25]. We performed three iterations of energy minimization for the set of wild-type residues in the viral spike protein bound with human ACE2 and mutant residues in the mink-CoV spike with mink ACE2. The relative binding energy (Δ*E*) are reported as the mean and standard deviation values across three replicates.

### Selective coefficient index

The selection coefficient index (*S*) of all SARS-CoV-2 codons was estimated by the FMutSel0 model in the program CODEML (PAML package) [26], The fitness parameter of the most common residues at each location is fixed to 0, while the other fitness parameters are limited to −20 < F < 20.

### Statistical analysis and mapping

Statistical analyses were performed using ANOVA followed by Turkey’s post hoc test, and the data were considered significantly different if the p-value was less than 0.05. ***p<0.001, **p<0.01, *p<0.05. The figures were mapped by the software PRISM GraphPad 5.0.

## Results

### Sequence and analysis of SARS-CoV-2 isolated from animals

As of Feb 2^nd^ 2021, more than 400 thousand SARS-CoV-2 genome sequences had been uploaded to the GISAID database. It is important to study the mutation rates and selective pressures on the SARS-CoV-2 genome during the spread of the epidemic. The results presented in Fig 1A show that the evolutionary entropy increased at specific sites in the whole genome of SARS-CoV-2, indicating substitution and selection capacity at these sites. In addition to humans, SARS-CoV-2 infects other animals (Fig 1B) and evolves in these animals. A phylogenetic tree was reconstructed based on animal-derived whole genome consensus sequences compared with the SARS-CoV-2 human isolate WIV04 (Fig 1C). Most SARS-CoV-2 clade isolates from the same animal clustered together, and the same clade contained sequences from all the mink regardless of their geographic region.

**Fig 1.**
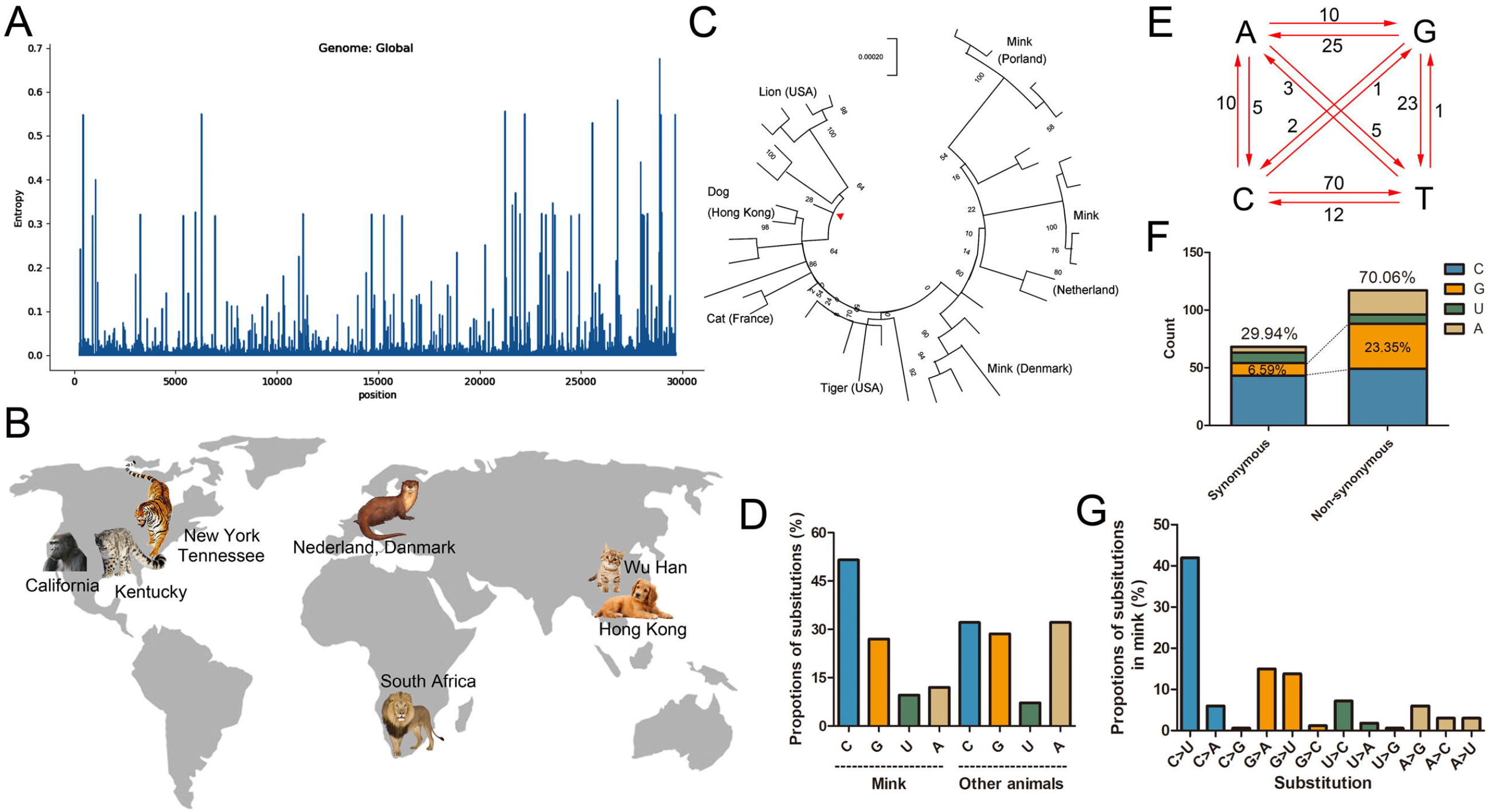
Composition and substitution analysis of SARS-CoV-2 isolated from animals. (A) The evolutionary entropy of specific sites on the SARS-CoV-2 genome in all the GISAID sequences on February 1, 2021. (B) The reported animals infected with SARS-CoV-2 and the defined transmission route from human to animal. (C) Phylogenetic tree using the maximum likelihood method and Tamura-Nei model performed by MEGA-X. The tree was provided with 500 bootstraps. (D) The proportions of uracil, guanine, thymine, and cytidine substitutions (nonsynonymous) in mink SARS-CoV-2 and other animals were separately counted. (E) Base pair changes observed in the mink SARS-CoV-2 genomes. All transitions and transversions were recorded and analyzed (see Supplementary Table S2). (F) The synonymous and nonsynonymous substitutions of mink-CoV were counted and analyzed. (G) The relative proportions of all transitions and transversions were separately analyzed.

The cluster of SARS-CoV-2 from mink (mink-CoV) has more substitutions compared to the reference sequence WIV04 (Supplementary Table S2), and the substitutions of cytidine in mink-CoV account for nearly 50% of the substitutions, while in other animals, cytidine accounts for only 30% of the substitutions (Fig 1D). The substitution of adenine in SARS-CoV-2 in other animals is threefold higher than that in mink-CoV. To track how the substitutions occurred in the mink-CoV genome, we recorded all the mutations in the mink-CoV genome in reference to the WIV04 genome. The results in Fig 1E & 1G show that the cytidine-to-uracil transition occurred more than 40% of the time and was eightfold higher than the uracil-to-cytidine substitution. Notably, the substitutions of guanine and adenine were more than threefold higher in nonsynonymous mutations than in synonymous mutations (Fig 1F).

### Mutational spectra of Spike protein in human and animal samples

The evolutionary entropy (Fig 2A) analysis revealed that most of the notable mutation pressures on the Spike protein occurred primarily in three relatively narrow domains, the N-terminal domain (NTD, green), receptor binding motif (RBM, purple), SD (pink), and CH and CD (blue) domains. The variation in the spike gene was evident when all the included sequences isolated from humans and animals were recorded in our study, which led to the identification of a number of highly variable residues, including L18F, A222V, S477N, P681H, S982A and D1118H (Fig 2B and 2C). A total of 12 relatively high-frequency amino acid variation sites were detected. Except for D614G, the substitution with the highest frequency was A222V. Notably, sequences in dogs (EPI 722380) had the most amino acid variant types in animals, and the dog strains EPI 730652 and EPI 699508, clustered together, contained the A222V and P681H mutations. C-to-U substitutions were scattered throughout the SARS-CoV-2 genome and accounted for 24.06% of the substitutions in the spike gene in of all epidemic strains analyzed as of February 2, 2021 (Fig 2D). Because of the widespread transmission of D614G (GAT>GGT), A-to-G substitutions accounted for 56.12% of all the monitored strains. The result of dN-dS indicates the natural selection for mutations in these specific sites in the *spike* gene (dN-dS>0 indicates positive selection, and dN-dS<0 indicates purification selection). Fig 2E shows that sites 222, 262, 439 and 614 were exposed to strong positive selection pressure, while positions 294, 413, 1018 and 1100 were subjected to purifying selection during evolution.

**Fig 2.**
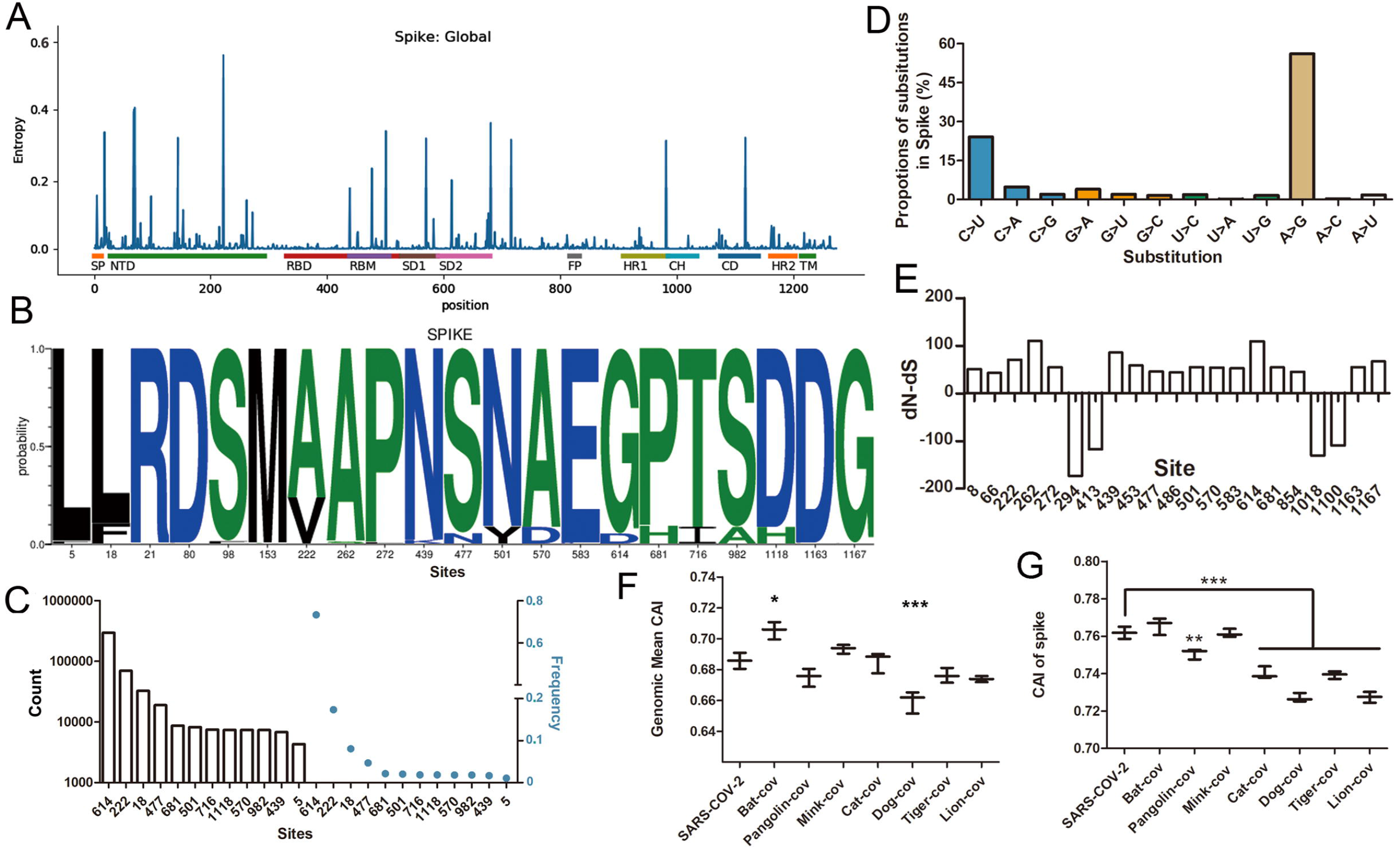
The mutation spectra of the spike protein and the selection pressure. (A) The evolutionary entropy of specific sites on the spike protein from all the GISAID sequences on February 1, 2021. (B) The WebLogo plots summarize the amino acid divergence of Spike sequences characterized in this study. The single letter amino acid (aa) code is used with the vertical height of the amino acid representing its prevalence at each position in the polypeptide (aa 18, 222, 477, 501, 570, 614, 982 and 1118 are indicated). (C) Total mutations of the Spike in the variants were recorded and counted after analysis by MEGA-X software. The frequency was calculated using the Datamonkey server. (D) The substitutions in the animal viral genome in this study were analyzed, including uracil, guanine, thymine, and cytidine as substituted with other bases. (E) The dN-dS value was calculated using the Datamonkey tool. (F) The genomic CAI value was calculated using SARS-CoV-2 sequences in humans and animals. Bat-CoV refers to RaTG13 and the corresponding host (bat), other animal-CoV means SARS-CoV-2 isolated from the indicted animals, the first SARS-CoV-2 refers the human host. (G) The CAI value of spike sequences in SARS-CoV-2 in humans and animals.

CAI was used to quantify the codon usage similarities between different coding sequences based on a reference set of highly expressed genes [27]. To clarify the optimization of SARS-CoV-2 in different hosts, we calculated the average CAI of the SARS-CoV-2 whole genome (Fig 2F) and spike region (Fig 2G). Interestingly, SARS-CoV-2 in bat hosts has a higher value of CAI relative to humans, while dogs had an obviously decreased CAI value compared to humans (Fig 2F). The bias of codon usage in the spike mutants are shown in Supplementary Table S3. Considering codon usage in the spike gene in different hosts, Fig 2G shows that pangolins, cats, dogs, tigers, and lions all had a lower CAI value than humans. These results indicated that SARS-CoV-2 optimized codon usage to adapt to the animals in which infection has been reported, but all of them showed a downward trend in adaptability relative to humans except for mink.

### Comparison of the receptors and binding affinity between humans and mammals

Recently, Wang et al. reported that the tyrosine-protein kinase receptor (UFO, also called AXL) is a candidate receptor for SARS-CoV-2 infection of the respiratory system [28]. Here, the interaction of spike with UFO was predicted using the ZDOCK sever (http://zdock.umassmed.edu/) after simulation with the structure of human and mink UFO. The results showed that the spike interacts with human and mink UFO through the amino acids Glu56, Glu59, His61, Glu70 and Glu85 (Fig 3B), which form electric charge attraction and hydrophobic interactions with residues K147, P251, D253 and N148 on spike. All these residues were located on the NTD of spike (Fig 3A). To distinguish the differentiation of receptor sequences between different animals and humans, the ACE2 and UFO amino acid sequences in humans, mink, ferrets, tigers, cats, and dogs were aligned (Fig 3C). The results showed that the critical mutations H34Y, L79H and G354R appear in mink and ferret ACE2 (Fig 3C upper), and the variations H61T, I68V and E85G are evident in the UFO sequences of all the animals except for tigers (Fig 3C lower). On the other hand, viral variation is another important factor that should also be considered when analyzing infection differences between animals and humans. Corresponding to the contact residues on the receptors, alignment of the viral sequence contacts of UFO and ACE2 on spike indicated that residues binding UFO are conserved (Fig 3D), while residues at site 453, which interact with those at position 34 in ACE2 (Fig 3E), showed a higher binding affinity for F453-Y34 in mink and ferrets than for Y453-H34 in humans (Fig 3F). The interaction of L486-T82 showed increased binding energy in mink and ferrets (Fig 3F). These variations indicate that the SARS-CoV-2 Spike shows a greater preference for binding the mink receptor ACE2 than human ACE2 after this mutation occurs.

**Fig 3.**
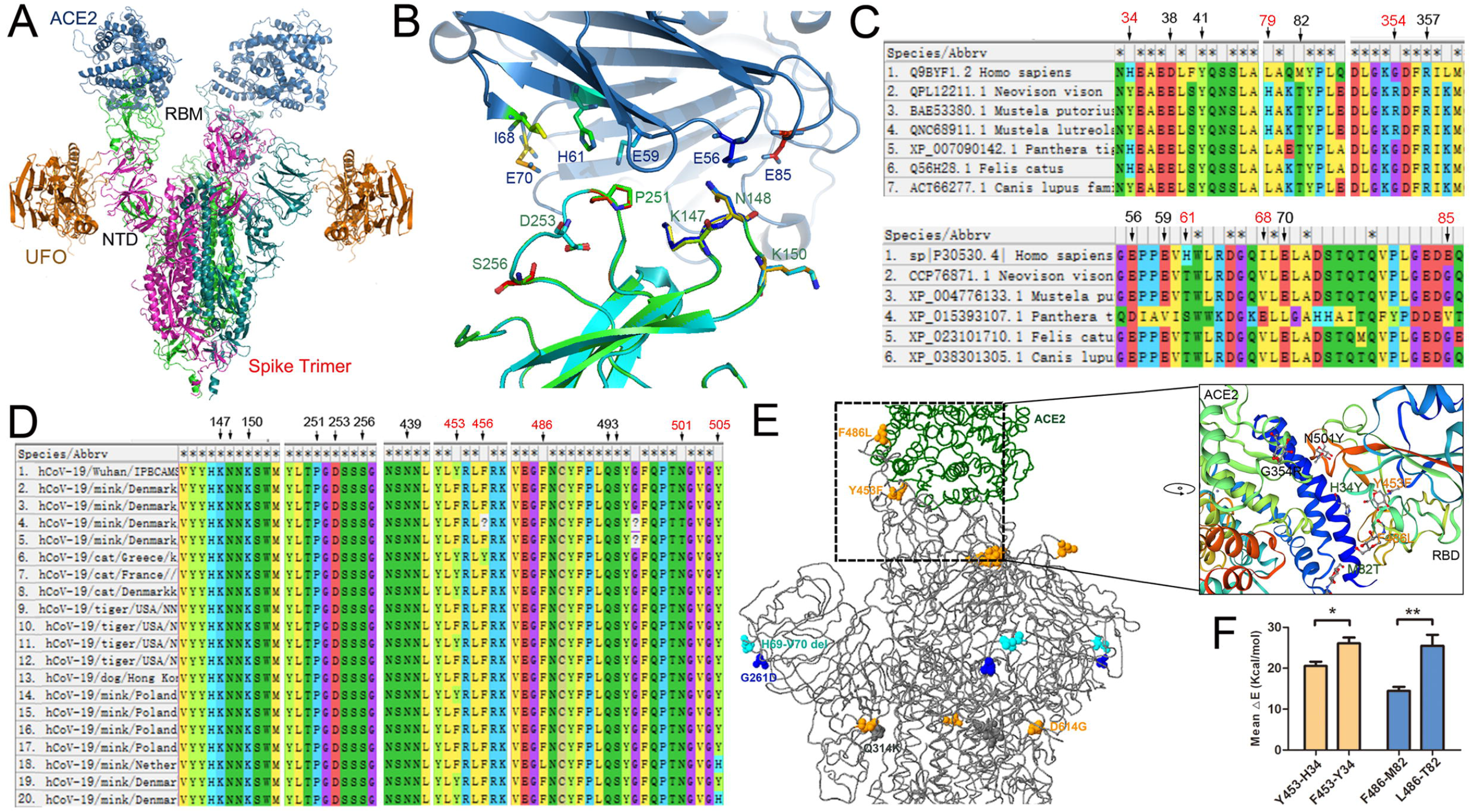
Receptors and binding analysis of animals and host adaptation. (A) Receptors ACE2 and UFO interacted with the SARS-CoV-2 Spike in different regions. (B) Human and mink UFO interacted with the SARS-CoV-2 NTD by hydrogen bonding. The human UFO is colored green, and the mink UFO is cyan. (C) Alignments of receptors ACE2 (upper) and UFO (lower) in humans and animals. The single letter amino acid (aa) that functions in the spatial interaction is indicated. (D) Alignment of the SARS-CoV-2 RBD sequences in humans and in animals reported to have been infected. The residues in contact with UFO (former) and ACE2 (latter) are indicated. (E) Results of the comparison of the spike structure from mink-CoV with the reference strain WIV04. Visualization of the changed residues within mink-CoV are shown as colored balls. (F) The binding free energy of the wild-type spike RBD with the human receptor and the mutant spike RBD with the mink receptor.

### Codon usage and mutations in the RBMs of spike proteins in mammals

Amino acid substitutions within the SARS-CoV-2 Spike RBM may have contributed to host adaption and cross-species transmission. N439K, S477N and N501Y were the most abundant variations throughout the RBM regions (Fig 4A and 4B). N439 does not bind directly with ACE2 but functions in the stabilization of the 498–505 loop [29], but the N439K substitution is absent in animal CoVs (Fig 3D). Previous computational analysis combined with entropy analysis of the spike (Fig 2A) showed that S477N may have decreased stability compared with the wild type [30]. Since human SARS-CoV-2 and mink-CoV do not show very different codon usage bias (Fig 2F) and because viral codon bias depends on the host, we compared the codon usage frequency of SARS-CoV-2 and SARS-CoV (Fig 4C), for which ferrets are common hosts. Because substitutions N501T (AAU>ACU) in mink and N501Y (AAU>UAU) in humans occurred nonsynonymously in the first and second positions and since these substitutions had a lower frequency than other noted substitutions (Fig 1G & 2D), further study on the relationship of these substitutions is needed. The results of selective coefficient index in Fig 4D show the differences of relative fitness in the SARS-CoV-2 codons, CGA and CGG have the high fitness score in all codon-specific estimates, and T (ACU) has a lower fitness than Y (UAU). In addition to the reported variation, other important mutations should also be considered in mink and human prevalent strains, such as Y505H (Fig 4E), which also affect binding with the ACE2 receptor and Histidine (CAU) has the similar codon fitness with Tyrosine (UAU) (Fig 4D).

**Fig 4.**
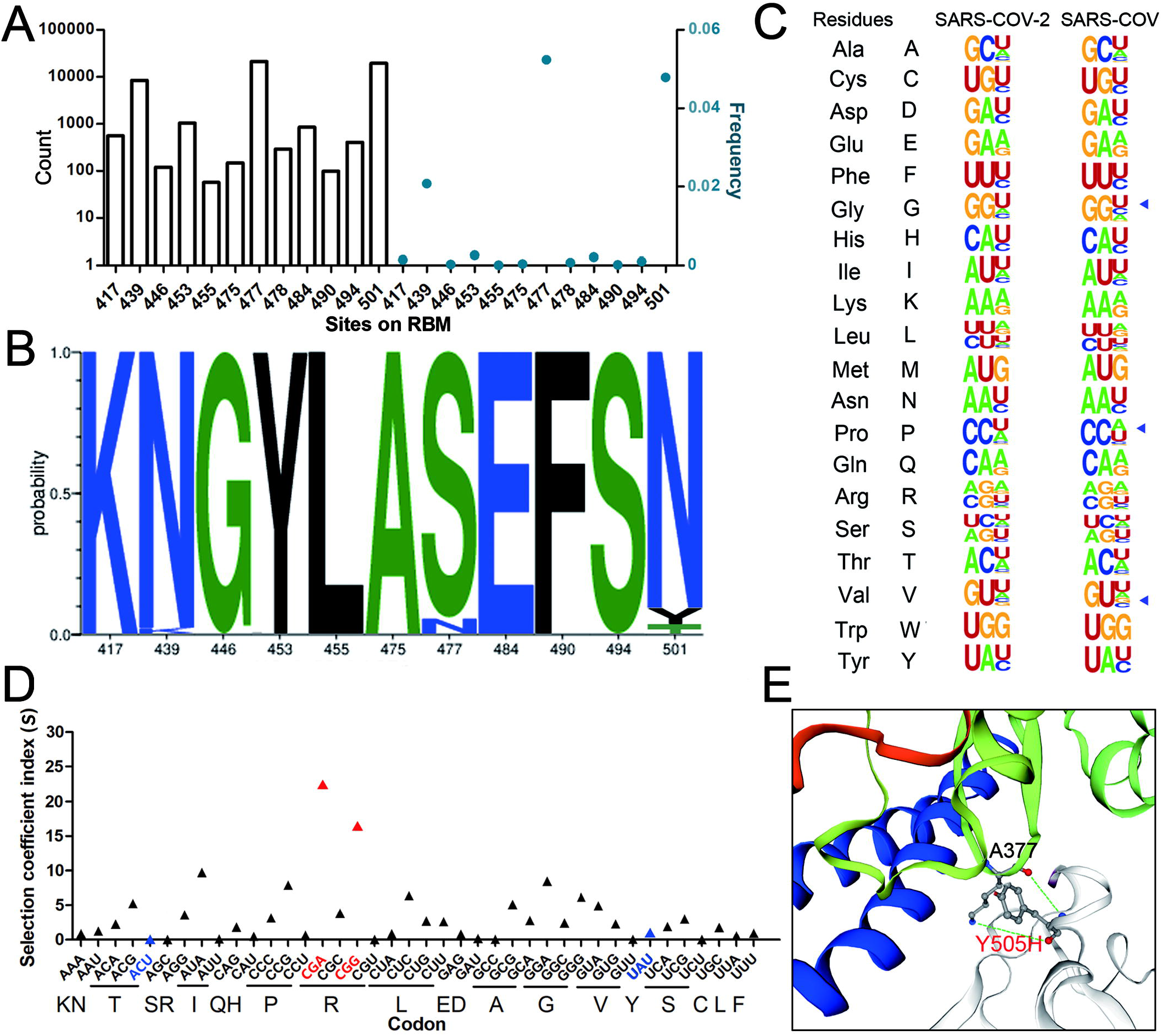
Codon usage and RBM mutations of the spike protein. (A) Total mutations of the RBM variants were recorded and counted after analysis by MEGA-X software. The frequency was calculated using the Datamonkey server. (B) The WebLogo plots summarize the amino acid divergence of RBM sequences from humans and mink. The single letter amino acid code is used with the vertical height of the amino acid representing its prevalence at each position in the polypeptide. (C) The synonymous codon usage bias of SARS-CoV-2 was produced by WebLogo (https://weblogo.berkeley.edu/logo.cgi), comparing the mink SARS-CoV-2 sequence MT396266 (GenBank ID) with SARS-CoV strain Toronto-2. (D) All codon-specific estimates of selective coefficient index were calculated, the indicated mutant codons of N501 were marked in blue, and the highest fitness codons were colored in red. (E) The variations Y505H in mink- or human-prevalent strains are also involved with binding to receptor ACE2.

In addition to the viral codon adaptation, mutation factors must be considered for virus prevalence. There was a lot lineages such as B.1.1.7, B.1.351, P.1 and the recently emerged lineage B.1.617 shared the same mutation sites Asp614 to Gly614 at spike (Fig 5A & 5B), and the B.1.351, P.1 possess the mutations E484K and N501Y which own the ability to escape natural and vaccine immunity system and have a broad prevalence in South Africa and Brazil (Fig 5D and 5E). The recently emerged lineage B.1.617 in India (Fig 5F) has two key mutations L452R and E484K at the same time, L452R confer resistance towards RBD-direct antibody and is characterized by a moderate increase in transmissibility. Increased surveillance needs to dare and will be crucial for control the epidemic and prevalence.

**Fig 5.**
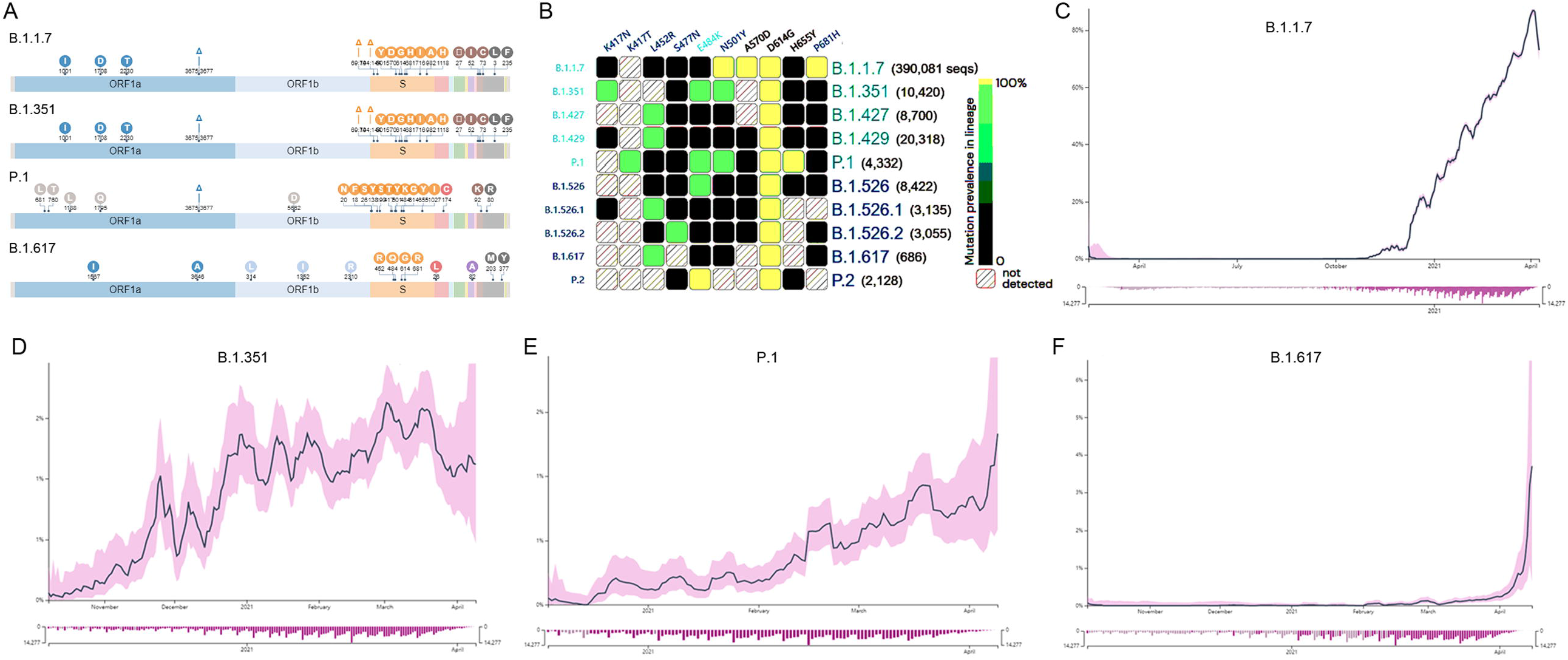
The prevalence of main lineages during the outbreak to April 2021. (A) The main characteristic lineages B.1.1.7, B.1.351, P.1 and B.1.617. (B) The mutations in RBD region of spike protein. (C) The prevalence of lineage B.1.1.7 in UK. (D) The prevalence of lineage B.1.351 in South Africa. (E) The prevalence of lineage P.1 in Brazil. (F) The prevalence of lineage B.1.617 in India.

## Discussion

Tracking animal variants arising from human contact or produced from animal bodies is an interesting topic and allows for better understanding of the evolutionary mechanism and selection fitness of SARS-CoV-2 in the host. Regardless of the probability of contact between different animals and SARS-CoV-2, the transmission of the virus between animals is inseparable from susceptibility and host adaptability. Mink were the first extensively farmed species to be affected by the COVID-19 epidemic, indicating that mustelids, including mink and ferrets, are more sensitive to SARS-CoV-2 than other animals [31]. Several mink farms in The Netherlands, Denmark, USA, and Spain all reported infection cases [32–35], indicating mink-to-mink and mink-to-human (Netherlands, Denmark) transmission. Other animals, including tigers and lions, are also susceptible to SARS-CoV-2 infection [36]. Hence, comparison of the susceptibility and the natural evolutionary pressure in different hosts for SARS-CoV-2 is meaningful and helpful for clarifying the host adaptation mechanisms and monitoring the epidemic.

Viral genes and genomes exhibit varying numbers of synonymous codons depending on the host [37]; hence, the codon usage bias of the virus has a strong relationship with its host. Studying the preferred synonymous codon usage and base substitutions helps to provide an understanding of the codon patterns of the virus in relation to their hosts and in relation to viral genome evolution. The convergence effect of virus codon preference on the host is widely recognized and is also one of the main natural selection forces for the coevolution of viruses and hosts [18]. In this study, we compared the codon bias of SARS-CoV-2 in mink with that of SARS-CoV in ferrets. Residues threonine (T) and tyrosine (Y) had similar codon biases in SARS-CoV-2 and SARS-CoV (Fig 4C), which both have the capability to infect mink and ferrets. The N501T variation mostly appeared in mink, while the N501Y mutation present only in humans cannot be explained from the perspective of codon bias and indicates that these two variations belong to two separate lineages.

The WebLogo diagram in Fig 4C shows that SARS coronaviruses preferentially have U- or A-ending codons. This is consistent with a previous report [38], and the G or C nucleotides in the third position of the preferred SARS-CoV-2 codons are not well represented. This feature may lead to an imbalance in the tRNA pool in infected cells, resulting in reduced host protein synthesis. The substitution rate of C-to-U was the highest in most of the reported sequences in animal species (Fig 1D). This may be because the surrounding context of cytidine in the sequence strongly influences the possibility of its mutation to U [39]. In the mink sequences, we observed an 8-fold increase in C-to-U substitution compared with the U-to-C substitution, which was higher than the reported 3.5-fold increase in mink [34], suggesting host adaptation of SARS-CoV-2 in mink over time and the ongoing outbreaks in multiple mink farms. In mink, the variations in G and A with nonsynonymous substitutions were higher than those with synonymous substitutions, which needs to be further analyzed. In addition, the sequences of other animal-CoVs are limited, such as those in the dogs and lions in the GISAID database, which is a limiting factor for comparison of base substitutions. CAI was used to measure the synonymous codon similarities between the virus and host coding sequences. For each animal source of the SARS-CoV-2 sequence, we calculated the average genome and *spike* gene values in the CAI (Fig 2F & 2G). Bat-CoV (RaTG13) and SARS-CoV-2 (from humans) had higher CAI values, which indicates that the viruses adapt to their hosts (bat and human) with optimized or preferred chosen codons, while the dog source of SARS-CoV-2 had lower CAI values, suggesting that SARS-CoV-2 adapts to dogs with random codons. This finding was consistent with the conclusion that, compared to dogs, humans are favored hosts for adaptation [40]. The whole genome or spike sequence in mink-CoV had a similar substitution level to human SARS-CoV-2, pointing to the ongoing adaptation of SARS-CoV-2 to the new host and using the preferred chosen codons.

The spike protein is critical for virus infection and host adaptation. We observed that three nonsynonymous mutations in the RBM domain, Y453F, F486L and N501T, independently emerged but were rarely observed in human lineages; these residues are directly involved in contact with the surface of the S-ACE2 complex and therefore are relevant to new-host adaptation. Other mutations within the RBM domain should also be monitored to prevent viral transmission and to further track the source. In addition to the mutation of the RBD, variations in the cell epitope of the spike protein should also be considered, and monitoring of the potential consequences of cell epitope variations in the process of viral transmission helps to adjust the vaccine strategy.

## Conclusions

Tracking animal variants arising from human contact or produced from animal bodies is an interesting topic and allows for better understanding of the evolutionary mechanism and selection fitness of SARS-CoV-2 in the host. Regardless of the probability of contact between different animals and SARS-CoV-2, the transmission of the virus between animals is inseparable from susceptibility and host adaptability. In this study, we systematically contrasted the position substitutions and codon usage of SARS-CoV-2 in human and animals, including dog, cat, lion, tiger and mink, showed the decreased adaptability of SARS-CoV-2 in animals relative to humans, except for mink. SARS-CoV-2 variant in mink showed a greater preference for binding with the mink receptor. This study focuses on the divergence of SARS-CoV-2 genome composition and codon usage in humans and animals, indicating possible natural selection and current host adaptation.

## DATA AVAILABILITY STATEMENT

All datasets presented in this study are included in the article/supplementary material.

## AUTHOR CONTRIBUTIONS

ZXL, DZ, RPY and FZ contributed to the design of experiments. ZXL, DZ, RPY, FZ, SRY, JJR and ZXL contributed to the conduction of experiments. ZXL, DZ, RPY, FZ, and JL contributed to the reagents. JL, WXD and ZXL contributed to the analyses of the data. LL and ZXL contributed to the writing the paper. LL contributed to the editing the paper.

## References

1. Zhu, N.; Zhang, D.; Wang, W.; Li, X.; Yang, B.; Song, J.; Zhao, X.; Huang, B.; Shi, W.; Lu, R.; et al. A Novel Coronavirus from Patients with Pneumonia in China, 2019. N Engl J Med 2020, doi:10.1056/NEJMoa2001017.

2. Zhou, P.; Yang, X.L.; Wang, X.G.; Hu, B.; Zhang, L.; Zhang, W.; Si, H.R.; Zhu, Y.; Li, B.; Huang, C.L.; et al. A pneumonia outbreak associated with a new coronavirus of probable bat origin. Nature 2020, doi:10.1038/s41586-020-2012-7.

3. Chan, J.F.; Kok, K.H.; Zhu, Z.; Chu, H.; To, K.K.; Yuan, S.; Yuen, K.Y. Genomic characterization of the 2019 novel human-pathogenic coronavirus isolated from a patient with atypical pneumonia after visiting Wuhan. Emerg Microbes Infect 2020, 9, 221–236, doi:10.1080/22221751.2020.1719902.

4. Letko, M.; Marzi, A.; Munster, V. Functional assessment of cell entry and receptor usage for SARS-CoV-2 and other lineage B betacoronaviruses. Nat Microbiol 2020, 5, 562–569, doi:10.1038/s41564-020-0688-y.

5. Conceicao, C.; Thakur, N.; Human, S.; Kelly, J.T.; Logan, L.; Bialy, D.; Bhat, S.; Stevenson-Leggett, P.; Zagrajek, A.K.; Hollinghurst, P.; et al. The SARS-CoV-2 Spike protein has a broad tropism for mammalian ACE2 proteins. PLoS biology 2020, 18, e3001016, doi:10.1371/journal.pbio.3001016.

6. Kim, Y.I.; Kim, S.G.; Kim, S.M.; Kim, E.H.; Park, S.J.; Yu, K.M.; Chang, J.H.; Kim, E.J.; Lee, S.; Casel, M.A.B.; et al. Infection and Rapid Transmission of SARS-CoV-2 in Ferrets. Cell host & microbe 2020, 27, 704–709 e702, doi:10.1016/j.chom.2020.03.023.

7. Shi, J.; Wen, Z.; Zhong, G.; Yang, H.; Wang, C.; Huang, B.; Liu, R.; He, X.; Shuai, L.; Sun, Z.; et al. Susceptibility of ferrets, cats, dogs, and other domesticated animals to SARS-coronavirus 2. Science 2020, 368, 1016–1020, doi:10.1126/science.abb7015.

8. Sit, T.H.C.; Brackman, C.J.; Ip, S.M.; Tam, K.W.S.; Law, P.Y.T.; To, E.M.W.; Yu, V.Y.T.; Sims, L.D.; Tsang, D.N.C.; Chu, D.K.W.; et al. Infection of dogs with SARS-CoV-2. Nature 2020, 586, 776–778, doi:10.1038/s41586-020-2334-5.

9. Zhang, Q.; Zhang, H.; Gao, J.; Huang, K.; Yang, Y.; Hui, X.; He, X.; Li, C.; Gong, W.; Zhang, Y.; et al. A serological survey of SARS-CoV-2 in cat in Wuhan. Emerg Microbes Infect 2020, 9, 2013–2019, doi:10.1080/22221751.2020.1817796.

10. Zhang, Q.; Zhang, H.; Huang, K.; Yang, Y.; Hui, X.; Gao, J.; He, X.; Li, C.; Gong, W.; Zhang, Y.; et al. SARS-CoV-2 neutralizing serum antibodies in cats: a serological investigation. bioRxiv 2020, 2020.2004.2001.021196, doi:10.1101/2020.04.01.021196.

11. McAloose, D.; Laverack, M.; Wang, L.; Killian, M.L.; Caserta, L.C.; Yuan, F.; Mitchell, P.K.; Queen, K.; Mauldin, M.R.; Cronk, B.D.; et al. From People to Panthera: Natural SARS-CoV-2 Infection in Tigers and Lions at the Bronx Zoo. mBio 2020, 11, doi:10.1128/mBio.02220-20.

12. Oreshkova, N.; Molenaar, R.J.; Vreman, S.; Harders, F.; Oude Munnink, B.B.; Hakze-van der Honing, R.W.; Gerhards, N.; Tolsma, P.; Bouwstra, R.; Sikkema, R.S.; et al. SARS-CoV-2 infection in farmed minks, the Netherlands, April and May 2020. Euro surveillance : bulletin Europeen sur les maladies transmissibles = European communicable disease bulletin 2020, 25, doi:10.2807/1560-7917.Es.2020.25.23.2001005.

13. Wang, L.; Mitchell, P.K.; Calle, P.P.; Bartlett, S.L.; McAloose, D.; Killian, M.L.; Yuan, F.; Fang, Y.; Goodman, L.B.; Fredrickson, R.; et al. Complete Genome Sequence of SARS-CoV-2 in a Tiger from a U.S. Zoological Collection. Microbiol Resour Announc 2020, 9, doi:10.1128/MRA.00468-20.

14. Daly, N. Several gorillas test positive for COVID-19 at California zoo—first in the world. 2021.

15. Andrew, S. Three snow leopards test positive for coronavirus, making it the sixth confirmed animal species. 2021.

16. Molenaar, R.J.; Vreman, S.; Hakze-van der Honing, R.W.; Zwart, R.; de Rond, J.; Weesendorp, E.; Smit, L.A.M.; Koopmans, M.; Bouwstra, R.; Stegeman, A.; et al. Clinical and Pathological Findings in SARS-CoV-2 Disease Outbreaks in Farmed Mink (Neovison vison). Vet Pathol 2020, 57, 653–657, doi:10.1177/0300985820943535.

17. Chaney, J.L.; Clark, P.L. Roles for Synonymous Codon Usage in Protein Biogenesis. Annu Rev Biophys 2015, 44, 143–166, doi:10.1146/annurev-biophys-060414-034333.

18. Bahir, I.; Fromer, M.; Prat, Y.; Linial, M. Viral adaptation to host: a proteome-based analysis of codon usage and amino acid preferences. Mol Syst Biol 2009, 5, 311, doi:10.1038/msb.2009.71.

19. Tamura, K.; Nei, M. Estimation of the number of nucleotide substitutions in the control region of mitochondrial DNA in humans and chimpanzees. Mol Biol Evol 1993, 10, 512–526, doi:10.1093/oxfordjournals.molbev.a040023.

20. Kumar, S.; Stecher, G.; Li, M.; Knyaz, C.; Tamura, K. MEGA X: Molecular Evolutionary Genetics Analysis across Computing Platforms. Mol Biol Evol 2018, 35, 1547–1549, doi:10.1093/molbev/msy096.

21. Dereeper, A.; Nicolas, S.; Le Cunff, L.; Bacilieri, R.; Doligez, A.; Peros, J.P.; Ruiz, M.; This, P. SNiPlay: a web-based tool for detection, management and analysis of SNPs. Application to grapevine diversity projects. BMC Bioinformatics 2011, 12, 134, doi:10.1186/1471-2105-12-134.

22. Sharp, P.M.; Li, W.H. The codon Adaptation Index--a measure of directional synonymous codon usage bias, and its potential applications. Nucleic acids research 1987, 15, 1281–1295, doi:10.1093/nar/15.3.1281.

23. Puigbo, P.; Bravo, I.G.; Garcia-Vallve, S. CAIcal: a combined set of tools to assess codon usage adaptation. Biol Direct 2008, 3, 38, doi:10.1186/1745-6150-3-38.

24. Xia, X. DAMBE5: a comprehensive software package for data analysis in molecular biology and evolution. Mol Biol Evol 2013, 30, 1720–1728, doi:10.1093/molbev/mst064.

25. Land, H.; Humble, M.S. YASARA: A Tool to Obtain Structural Guidance in Biocatalytic Investigations. Methods Mol Biol 2018, 1685, 43–67, doi:10.1007/978-1-4939-7366-8_4.

26. Yang, Z.; Nielsen, R. Mutation-selection models of codon substitution and their use to estimate selective strengths on codon usage. Mol Biol Evol 2008, 25, 568–579, doi:10.1093/molbev/msm284.

27. Henry, I.; Sharp, P.M. Predicting gene expression level from codon usage bias. Mol Biol Evol 2007, 24, 10–12, doi:10.1093/molbev/msl148.

28. Wang, S.; Qiu, Z.; Hou, Y.; Deng, X.; Xu, W.; Zheng, T.; Wu, P.; Xie, S.; Bian, W.; Zhang, C.; et al. AXL is a candidate receptor for SARS-CoV-2 that promotes infection of pulmonary and bronchial epithelial cells. Cell Res 2021, 31, 126–140, doi:10.1038/s41422-020-00460-y.

29. Li, F.; Li, W.; Farzan, M.; Harrison, S.C. Structure of SARS coronavirus spike receptor-binding domain complexed with receptor. Science 2005, 309, 1864–1868, doi:10.1126/science.1116480.

30. Mathavan, S.; Kumar, S. Evaluation of the Effect of D614g, N501y and S477n Mutation in Sars-Cov-2 through Computational Approach. 2020, doi:10.20944/preprints202012.0710.v1.

31. Manes, C.; Gollakner, R.; Capua, I. Could Mustelids spur COVID-19 into a panzootic? Vet Ital 2020, 56, 65–66, doi:10.12834/VetIt.2375.13627.1.

32. Jo, W.K.; de Oliveira-Filho, E.F.; Rasche, A.; Greenwood, A.D.; Osterrieder, K.; Drexler, J.F. Potential zoonotic sources of SARS-CoV-2 infections. Transbound Emerg Dis 2020, doi:10.1111/tbed.13872.

33. Opriessnig, T.; Huang, Y.W. Further information on possible animal sources for human COVID-19. Xenotransplantation 2020, 27, e12651, doi:10.1111/xen.12651.

34. Oude Munnink, B.B.; Sikkema, R.S.; Nieuwenhuijse, D.F.; Molenaar, R.J.; Munger, E.; Molenkamp, R.; van der Spek, A.; Tolsma, P.; Rietveld, A.; Brouwer, M.; et al. Transmission of SARS-CoV-2 on mink farms between humans and mink and back to humans. Science 2021, 371, 172–177, doi:10.1126/science.abe5901.

35. Mahdy, M.A.A.; Younis, W.; Ewaida, Z. An Overview of SARS-CoV-2 and Animal Infection. Front Vet Sci 2020, 7, 596391, doi:10.3389/fvets.2020.596391.

36. Csiszar, A.; Jakab, F.; Valencak, T.G.; Lanszki, Z.; Toth, G.E.; Kemenesi, G.; Tarantini, S.; Fazekas-Pongor, V.; Ungvari, Z. Companion animals likely do not spread COVID-19 but may get infected themselves. Geroscience 2020, 42, 1229–1236, doi:10.1007/s11357-020-00248-3.

37. Lloyd, A.T.; Sharp, P.M. Evolution of codon usage patterns: the extent and nature of divergence between Candida albicans and Saccharomyces cerevisiae. Nucleic acids research 1992, 20, 5289–5295, doi:10.1093/nar/20.20.5289.

38. Alonso, A.M.; Diambra, L. SARS-CoV-2 Codon Usage Bias Downregulates Host Expressed Genes With Similar Codon Usage. Front Cell Dev Biol 2020, 8, 831, doi:10.3389/fcell.2020.00831.

39. Simmonds, P. Rampant C-->U Hypermutation in the Genomes of SARS-CoV-2 and Other Coronaviruses: Causes and Consequences for Their Short- and Long-Term Evolutionary Trajectories. mSphere 2020, 5, doi:10.1128/mSphere.00408-20.

40. Dutta, R.; Buragohain, L.; Borah, P. Analysis of codon usage of severe acute respiratory syndrome corona virus 2 (SARS-CoV-2) and its adaptability in dog. Virus research 2020, 288, 198113, doi:10.1016/j.virusres.2020.198113.

