## Supplemental Table 3 for "Substitutions and codon usage in SARS-CoV-2 in mammals indicate natural selection and host adaptation" (from another version of this preprint: version 1, 2021-04-05)

Table S3. The codon bias of residues on the spike and RBM

| Position within Spike | Codon | Substitution | Position within Spike | Codon | Substitution |
| --- | --- | --- | --- | --- | --- |
| 5 | CUU | UUU | 501 | AAU | UAU |
| 18 | CUU | UUU | 570 | GCU | GAU |
| 21 | AGA | AUU | 583 | GAG | GAC |
| 80 | AGU | UAU | 614 | GAU | GGU |
| 98 | UCU | UUU | 681 | CCU | CAU |
| 153 | AUG | ACG | 716 | ACA | AUA |
| 222 | GCU | GUU | 982 | UCA | GCA |
| 262 | GCU | UCU | 1118 | GAC | CAC |
| 272 | CCU | CUU | 1163 | GAU | UAU |
| 439 | AAC | AAA | 1167 | GGU | GUU |
